# Swimmer’s itch control: timely waterfowl brood relocation significantly reduces an avian schistosome population and human cases on recreational lakes

**DOI:** 10.1101/2023.07.10.548414

**Authors:** Curtis L. Blankespoor, Harvey D. Blankespoor, Randall J. DeJong

## Abstract

Swimmer’s itch (SI) is a dermatitis in humans caused by cercariae of avian and mammalian schistosomes which emerge from infected snails on a daily basis. Mitigation methods for SI have long been sought with little success. Copper sulfate application to the water to kill the snail hosts is the historically employed method, but is localized, temporary, and harmful to many aquatic species. Here, we test an alternative method to control *Trichobilharzia stagnicolae*, a species well-known to cause SI in northern Michigan and elsewhere in North America. Summer relocation of broods of the only known vertebrate host, common merganser (*Mergus merganser),* greatly reduced snail infection prevalence the following year on two large, geographically separated lakes in northern Michigan. Subsequent years of host relocation achieved and maintained snail infection prevalence at ∼0.05%, more than an order of magnitude lower than pre-intervention. A Before – After – Control – Intervention (BACI) study design using multiple-year snail infection data from two intervention lakes and three control lakes demonstrates that substantial lake-wide reduction of an avian schistosome can be achieved and is not due to natural fluctuations in the parasite populations. The relevance of reducing snail infection prevalence is demonstrated by a large seven-year data set of SI incidence in swimmers at a high-use beach, which showed a remarkable reduction in SI cases in two successive years after relocation began. In addition, data from another Michigan lake where vertebrate-host based intervention occurred in the 1980’s is analyzed statistically and shows a remarkably similar pattern of reduction in snail infection prevalence. Together, these results demonstrate a highly effective SI mitigation strategy that avoids the use of environmentally suspect chemicals and removes incentive for lethal host removal. Biologically, the results strongly suggest that *T. stagnicolae* is reliant on the yearly hatch of ducklings to maintain populations at high levels on a lake and that the role of migratory hosts in the spring and fall is much less significant.

## Introduction

Swimmer’s itch, or cercarial dermatitis, is caused by parasitic flatworms in the family Schistosomatidae, which require both snail intermediate hosts and avian or mammal definitive hosts to complete their life cycles. Adult worms live in avian or mammal hosts, producing embryonated eggs which exit the host, typically in the feces. Upon contact with water, the eggs hatch quickly into free-living miracidia, which seek out the appropriate host snail species, penetrate the integument, and begin to develop into sporocysts. After a few weeks, sporocysts asexually produce cercariae, free-living stages that emerge from the snail. A single infected snail can produce up to several thousand cercariae per day. If cercariae encounter a suitable vertebrate host, they penetrate the skin or other epithelial surfaces (e.g., mouth or throat), then migrate in the host and enter the vasculature, where they mature into male and female worms that reproduce sexually.

When cercariae of nonhuman schistosomes encounter human skin, they can penetrate, but ultimately do not survive. Reddened, raised areas called papules usually result at the points of entry, and can itch intensely for up to a week or more. Cercariae are planktonic and can be dispersed or concentrated by wind and current, and persons spending time in the water can contract hundreds of papules. Papules develop in most people even on the first exposure [1–3], but subsequent exposures can also elicit stronger and more intense reactions [2]. The resulting condition can range from mild annoyance to intense discomfort requiring medical attention [4]. In addition, there is recognition in the literature of a potential public health concern because some species may briefly survive beyond the skin and migrate within incorrect hosts [5–9].

Swimmer’s itch (SI) has been described from a wide range of latitudes and has been linked to multiple schistosome genera, although *Trichobilharzia* is the most diverse and is the most often linked to SI outbreaks in Europe and North America [10–15]. In the northern part of the Michigan lower peninsula, where the link with avian schistosomes was first made [16], SI is a common issue on many inland lakes. Although multiple schistosome species can be present in a given lake, older and recent studies show *T. stagnicolae* to be the dominant schistosome on multiple lakes in northern Michigan [17–23], especially on oligotrophic lakes. In addition, a recent study demonstrated that *T. stagnicolae* is many times better at penetrating human skin [24] relative to another species present in Michigan. *Trichobilharzia stagnicolae* utilizes *Stagnicola* snails (Lymnaeidae) as intermediate hosts, and common mergansers (*Mergus merganser*) as definitive hosts. When surveyed, adult and hatch-year common mergansers have been found to be very frequently infected with schistosomes [13,14,25,26], with high numbers of parasite eggs excreted. Molecular analyses of eggs passed from common mergansers in Michigan [17,27] have confirmed that they are infected with *T. stagnicolae* and *T. physellae*, with the latter schistosome species requiring physid snail hosts that are present but usually less common on these lakes [28].

The presence of SI in places where tourism is strongly connected to recreational water use may lead to significant economic losses [10,14,15,29,30]. Multiple methods to mitigate SI have been proposed or attempted (see Soldánová et al. [15] for a thorough review) but no one solution has been recognized in the literature as reliably effective [10,12,15]. The most common method implemented continues to be copper sulfate as a molluscicide which can reduce but not eliminate snail populations in a treated area [25,31,32]. However, cercariae from outside the application area may drift into the treated area [32], and copper sulfate use is restricted in many U.S. states because of toxicity to other invertebrates, algae, and fish.

Using a Before – After – Control – Impact/Intervention (BACI) study design [33–35], we tested whether relocating common merganser broods is an effective method to reduce the presence of *T. stagnicolae* and the incidence of SI on two Michigan lakes. We predicted that reductions in snail infection prevalence and SI incidence would be observed in the immediate year after relocation and would be sustained in subsequent years with continued brood relocation. Snail infection data from very large numbers of snails collected before and after the intervention and a dataset representing multiple years of recorded SI cases are used to assess these predictions. The analyses are strengthened by concurrent snail infection data from control lakes where relocation efforts did not occur and by historical snail infection data previously published.

## Methods

### Intervention Lakes

All lakes in this study are located in the northern half of the Michigan lower peninsula. Common merganser broods were relocated from Higgins Lake (Roscommon County) starting in 2015. Since it is the young snails that become infected in the summer, overwinter, and carry *T. stagnicolae* infections into the next summer, 2015 snail data are reflective of conditions ‘Before’ brood relocation could have an effect. Snail data from 2016 and onwards is reflective of ‘After’ brood relocation at Higgins Lake. Common merganser brood relocation on Crystal Lake (Benzie County) began in 2017, making snail data from 2018 and onwards ‘After’ the effects of Crystal Lake brood relocation. Both intervention lakes are oligotrophic with a predominately sandy substrate. The two intervention lakes are separated by 105 kilometers.

### Control Lakes

Big Glen Lake (Leelanau County), North Lake Leelanau (Leelanau County), and Douglas Lake (Cheboygan County) served as control lakes as common mergansers broods were present but no brood relocation was conducted prior to snail sampling. Big Glen Lake and North Lake Leelanau are similarly oligotrophic with sandy substrates, whereas Douglas Lake is considered ‘mesotrophic with some oligotrophic features’ [36], with extensive areas of sand. The three control lakes are 94-120 km from Higgins Lake and 24-150 km from Crystal Lake. The original study design intended for Big Glen Lake and North Lake Leelanau to be control lakes throughout the study, but early success of the relocation program on Higgins Lake resulted in these lakes beginning their own relocation programs, implemented and assessed by another group [28]. Since data from these lakes after 2017 could no longer serve as control data for our study, snail data was no longer collected from these lakes.

### Snail and schistosome diversity

Though the control lakes have somewhat higher snail diversity, the dominant snail on all five of these lakes is *Stagnicola emarginata* (=*Lymnaea catascopium* (18,22,25,28, personal observations)) which have lifespans of up to 15 months, from spring/early summer to late summer of the following year, (36–38, personal observations). The dominant schistosome on these lakes is *T. stagnicolae* [17,18,22,23], although other snails are present that can be intermediate hosts to schistosomes of other waterfowl such as Canada goose [24,40] and mallard [17]. However, the contributions of these other schistosome species to SI cases is likely minor compared to *T. stagnicolae*, as most are less frequently detected [17,23] and their snail hosts are lower in abundance. *Physell*a (=*Physa*) *parkeri* and other physid snails are present at low abundance at all of the lakes in this study excep Glen Lake, where *P. parkeri* is abundant [28]. *Physella* is host to *T. physellae,* a parasite of mallards and common mergansers that is present at all five lakes but at lower frequency and abundance [17,23]. *Planorbella* (=*Helisoma*) *trivolvis* is present on these lakes (very low to moderate abundance) and is host to a recently discovered, unnamed schistosome that uses Canada goose as vertebrate host. While *P. trivolvis* infected with this species has been found in Big Glen Lake and this parasite is widespread in Michigan and elsewhere [23,24,40], we recently demonstrated that this schistosome has a significantly lower propensity to penetrate human skin compared to *T. stagnicolae* [24]. Finally, *Gyraulus* spp. are small snails that are typically localized to particular microhabitats on these lakes and are not common. Schistosomes that use *Gyraulus* have only been detected on one lake in northern Michigan [17], though we assume they may be present in low abundance.

### Common merganser populations and brood relocation

All trapping, banding, and relocation activities by the authors were conducted with permission from the USFWS (Scientific Collecting permit MB54823B), the USGS (Bird Banding Master permit 24094), and the Michigan Department of Natural Resources (MI-DNR) (Scientific Collecting permit SC1543). IACUC oversight and approval was provided by the University of Michigan (PRO00010637).

On intervention lakes, all common merganser brooding hens and their ducklings were captured using a specially designed drive trap. Hens were banded with USFWS bands, and in 2017-2019, most ducklings were marked with web tags. All captured birds were transported in specialized animal transport cages to Michigan Department of Natural Resources (MI-DNR) approved relocation sites on Lake Michigan or Lake Huron, where common mergansers naturally breed in large numbers. Due to strong wave action, suitable host snails are notably absent from most of their shorelines and SI cases are rarely reported from Lake Michigan, Lake Huron, and the other three Great Lakes. Total numbers of brooding hens and ducklings relocated are provided in Table 1. We were also able to verify that common merganser broods were present on control lakes in the relevant years before snail work was performed, though counts were not obtained (Table 1).

**Table 1.** Presence and relocation data for common merganser broods on intervention and control lakes.

| Lake, Year | Number of common<br>merganser broods<br>(number of ducklings) | Number of common<br>merganser broods relocated<br>(number of ducklings) |
| --- | --- | --- |
| Big Glen Lake |  |  |
| 2014 | Present | No relocation |
| 2015 | Present | No relocation |
| 2016 | Present | No relocation |
| Crystal Lake |  |  |
| 2016 | At least 5 broods | No relocation |
| 2017 | 10 (116) | 10 (116) |
| 2018 | 14 (130) | 14 (130) |
| 2019 | 10 (68) | 10 (68) |
| Douglas Lake |  |  |
| 2014 | Present | No relocation |
| 2015 | Present | No relocation |
| 2018 | Present | No relocation |
| Higgins Lake |  |  |
| 2014 | Present | No relocation |
| 2015 | 9 (79) | 9 (79) |
| 2016 | 6 (47) | 6 (47) |
| 2017 | 4 (54) | 4 (54) |
| 2018 | 2 (16) | 2 (16) |
| 2019 | 2 (15) | 2 (15) |
| North Lake Leelanau |  |  |
| 2015 | Present | No relocation |

Second-year common mergansers do not breed, but commonly are present on inland lakes during which time females spend time prospecting for nest sites. Summer populations of second-year individuals were observed on both intervention lakes every year. They were counted during 1-4 bird surveys (slowly driving a boat once around the entire lakeshore to count all waterfowl) per year in June and July; for Higgins Lake the median was 8 second-year individuals (15 surveys, 2015-2020, count range 3-19 per survey), for Crystal Lake the median was 9 (4 surveys, 2016-2018, range 4-12 per survey).

Lethal take of common mergansers was not part of the study design of this project. Unfortunately, Gerrish township officials on Higgins Lake (unassociated with this study or the authors) were permitted to shoot 23 adult common mergansers in the spring migratory season (mostly in April) in 2015 and 2016. However, this represents only a portion of the Higgins Lake spring migratory population as inland lakes in Michigan have migrant populations that are several times larger, with most spring migrants continuing north to other breeding sites. Sightings of groups of 10-100+ common mergansers are common in spring and fall at both Crystal Lake and Higgins Lake (personal observations, eBird). Additionally, five individuals that had been wounded in the spring by Gerrish officials were lethally taken in summer 2015, along with four second-year individuals under our scientific collecting permit. An additional individual wounded by Gerrish officials was taken in 2016 at Higgins Lake, as was an individual observed to be wounded (unknown cause) in 2017 at Crystal Lake. Finally, a limited number of adult common mergansers were lethally taken for other scientific purposes under our scientific collecting permit in 2017 (5, Higgins Lake; 9, Crystal Lake) and 2018 (3, Crystal Lake), but in all cases there remained more breeding and second-year adults than were eliminated.

### Snail collection and cercarial shedding

*Stagnicola emarginata* snails were collected by wading and/or by snorkeling in depths up to 3 meters. Modified scoops, with mesh allowing sand to pass through, were used to collect snails off the lake bottom. Snails were housed in 5-gallon buckets filled with lake water, cooled with fresh lake water as needed, and covered overnight in the laboratory. The next morning, all snails were individually isolated in cups or in the wells of tissue culture trays with well water [41] and exposed to natural and artificial light for a minimum of 1.5 hours. Each snail, and any emerging cercariae, were observed under a light microscope. All schistosome infections were recorded, and for most site collections (305 of 314, >97%), all other trematode infections were also recorded by cercarial type. Recording the emergence of other trematodes from collected samples provides verification that snail isolation and observation methods were effective. Snail densities at most sites appeared to remain relatively consistent throughout the study, although noticeable shifts in snail beds toward shallower or deeper water occurred at some sites. Similar shifts have been described to occur within short time frames (a few weeks) for *S. emarginata* at Higgins Lake [39] and for multiple species of lymnaeids, including *S. emarginata,* at Douglas Lake [37].

In total, 58524 snails were collected and examined for trematode infections over six years (2015-2020). Numbers of snails examined and found to be infected are summarized in Table 3. Snail sampling efforts targeted and usually achieved a goal of 200 snails per site per collection date (248 of 314 site collections (79%) included 200 or more snails), though the number collected varied with snail abundance (41 site collections (13%) included 100-200 snails; 25 site collections (8%) included less than 100). Ten sites were sampled for each of the two intervention lakes, Crystal Lake and Higgins Lake, while fewer sites were sampled on the control lakes (Big Glen Lake, Douglas Lake, North Lake Leelanau; Table 3). Sampling Crystal Lake, Higgins Lake, and North Lake Leelanau sites 4-5 times in 2015-2017 enabled us to confirm that snail infections are readily detectable from the beginning of June to early August, when many snails begin to die [39], and that the highest prevalences are found from late June to late July (S1 Fig), consistent with peak detection in other studies [28,42]. Thereafter, we sampled snails on intervention lakes with a single collection of each site in mid-July, with a target of 2000 snails per lake.

### Historical snail infection data

*Stagnicola* snails from the lakes in this study (see lake descriptions above) were frequently sampled in the past and we utilize snail infection data from five older studies to enrich our analyses. The first data set is from an eight-year study (1983-1990) of Big Glen Lake before and after similar SI reduction methods were implemented [25]. Another set of data [18] includes Higgins Lake and North Lake Leelanau in years (1998-2001) in which definitive hosts were not disturbed [43]. Finally, Keas and Blankespoor [22] presented data from North Lake Leelanau (1990) and Douglas Lake (1994), and compared Douglas Lake data from two older studies [44,45]. All of these historical prevalence data are presented in Table 2.

**Table 2.** Historical prevalences of *T. stagnicolae* in *Stagnicola emarginata* from lakes in this study..

| Lake, year | Number of snails examined | Percent *T. stagnicola* infection (number of snails infected) | Data source |
| --- | --- | --- | --- |
| Big Glen Lake |  |  |  |
| 1983 | 15600 | 1.20% (187) | [25] |
| 1984 | 14252 | 1.19% (170) | [25] |
| 1985 | 4219 | 2.01% (85) | [25] |
| 1986 | 2498 | 0.36% (9) | [25] |
| 1987 | 10500 | 0.40% (42) | [25] |
| 1988 | 6400 | 0.09% (6) | [25] |
| 1989 | 6100 | 0.11% (7) | [25] |
| 1990 | 6240 | 0.06% (4) | [25] |
| Douglas Lake |  |  |  |
| 1935-36 | 4784 | 4.74% (227) | [44] |
| 1957 | 463 | 1.08% (5) | [45] |
| 1994 | 550 | 2.91% (16) | [22] |
| Higgins Lake |  |  |  |
| 1998 | 5866 | 1.06% (62) | [18] |
| 1999 | 8862 | 0.84% (74) | [18] |
| 2000 | 10633 | 0.56% (60) | [18] |
| 2001 | 5844 | 0.78% (46) | [18] |
| North Lake Leelanau |  |  |  |
| 1990 | 13213 | 1.25% (165) | [22] |
| 1999 | 2274 | 0.75% (17) | [18] |
| 2000 | 2636 | 1.06% (28) | [18] |
| 2001 | 2901 | 0.14% (4) | [18] |

Years at Big Glen Lake after intervention are marked with gray shading

### Crystal Lake CSA beach data

A large beachfront on the southwestern shore of Crystal Lake is owned and operated by the Congregational Summer Assembly (CSA), and an extensive offshore, members-only area is cordoned for recreational swimming and swimming lessons. Lifeguards at CSA collected data daily for 7 weeks from late June to mid-August from 2013-2019. Swimmers self-reported SI cases to lifeguards and daily tallies were recorded. Awareness about SI and motivation to report was high due to educational efforts within the CSA and a desire to record the severity of the SI problem. Lifeguards also counted the number of swimmers each session. The morning session ran from 0800-1100 h and the counts were performed at ∼1000 h. The afternoon session occurred from 1300 to 1700 h, and counts were taken at ∼1600 h. The beach is open at all times, but lifeguards are only on duty during the two sessions. Part of this data set (years 2013-2016, prior to any merganser relocation) was reported in an earlier publication that examined the role of environmental variables in predicting SI risk [46]. The additional years of data reported here provide another year of swimming data before the effects of common merganser relocation (2017), and two years of swimming data after the relocation efforts began (2018, 2019). Data were not collected in 2020 due to the COVID-19 pandemic. The data represent totals of 354 swimming days, 36838 swimmers, and 1244 SI cases, and the data are approved for research use (Institutional Review Board of Calvin University, 21-013).

### Data analyses

We fit a mixed-effects logistic regression model to predict the probability of collecting snails infected with *T. stagnicolae* as a function of control vs. intervention lake and before vs. after common merganser relocation with the random effect of collection site. Because mitigation efforts targeting definitive hosts were begun at different times on Higgins Lake (2015) and Crystal Lake (2017), separate models were fit for each intervention lake compared to the appropriate years of data for control lakes. A separate model was also fit for Big Glen Lake as the intervention lake using historical data from other lakes for control lakes. All models were fitted with R version 4.2.0 using package glmmTMB [47,48] and all R markdown files for these analyses are available.

For the CSA beach, onshore wind directions of N and NE were found by Sckrabulis et al., [46] to be highly predictive of SI cases in the 2013-2016 data. In the larger data set analyzed here (2013-2019), 83.8% of the SI cases occurred on days with N or NE winds (17.5% and 66.2% of cases, respectively). The frequency of days with NNE winds was similar before and after intervention (32.2% before, 30.6% after). We fit a logistic regression model, using the quasi-binomial family to account for overdispersion, to predict the probability of a swimmer getting SI at CSA beach as a function of NNE wind directions and before vs. after common merganser relocation. This model was also fitted with R version 4.2.0 [48] and R markdown files are available.

## Results

### Snail infection data

Lake-wide percentages of snails infected with *T. stagnicolae* decreased significantly following annual common merganser relocation at Higgins Lake and Crystal Lake (Table 3 and Fig 1; X^2^= 121.6, p < 2 x 10^-16^, Higgins Lake; X^2^= 4.3, p = 0.038, Crystal Lake). Reductions in snail infection rate were seen immediately in the first year with a ∼3-fold reduction on Crystal Lake, and ∼10-fold reduction on Higgins Lake (Table 3). Moreover, these reductions were sustained for every year measured after relocation began and reached lows of ∼0.05% on both lakes, for a total reduction of greater than 10-fold on Crystal Lake and more than 50-fold on Higgins Lake (Table 3). The effects were lake-wide with decreases at every sampling site. A second analysis controlled for time of collection by restricting comparisons to yearly collections in the first half of July (Table 4). Statistical models fit on these July data also showed that even with these much smaller datasets the decreases in snail infection percentage on impact lakes were substantial (model values X^2^= 56.8, p = 4.9 x 10^-14^, Higgins Lake; X^2^= 13.0, p = 0.0003, Crystal Lake). A similar outcome was found for Big Glen Lake, with lake-wide percentages of snails infected with *T. stagnicolae* substantially lower as a result of intervention and remarkably similar to those found in our study (Table 3 and Fig 2; X^2^=8.2, p = 0.0042).

**Table 3.** Collection and infection data for *Stagnicola emarginata* snails examined in this study.

| Lake, Year | Number of sites | Number of collections per site | Number of snails examined | Percent *T. stagnicola* infection (number of infected snails) | Percent other trematode infections (number of infected snails) |
| --- | --- | --- | --- | --- | --- |
| Big Glen Lake |  |  |  |  |  |
| 2015 | 4 | 2 | 1597 | 0.38% (6) | 3.01% (48) |
| 2016 | 4 | 2 | 1479 | 0.20% (3) | 7.57% (112) |
| 2017 | 4 | 2 | 1761 | 3.01% (53) | 6.13% (108) |
| Crystal Lake |  |  |  |  |  |
| 2016 | 10 | 5 | 10212 | 0.79% (81) | 6.11% (624) |
| 2018 | 10 | 1 | 2112 | 0.28% (6) | 4.36% (92) |
| 2020 | 10 | 1 | 2211 | 0.05% (1) | 3.21% (71) |
| Douglas Lake |  |  |  |  |  |
| 2015 | 3 | 1-2 | 422 | 4.74% (20) | ---- |
| 2016 | 3 | 1-2 | 379 | 0.79% (3) | ---- |
| 2019 | 2 | 1 | 807 | 0.87% (7) | 7.43% (60) |
| Higgins Lake |  |  |  |  |  |
| 2015 | 10 | 5 | 7734 | 3.01% (233) | 8.68% (671) |
| 2016 | 10 | 5 | 9948 | 0.28% (28) | 6.13% (610) |
| 2017 | 10 | 4 | 8358 | 0.05% (4) | 3.31% (277) |
| 2019 | 10 | 1 | 1383 | 0.07% (1) | 8.68% (120) |
| 2020 | 10 | 1 | 2063 | 0.00% (0) | 9.79% (202) |
| North Lake Leelanau |  |  |  |  |  |
| 2016 | 8 | 5 | 8058 | 0.66% (53) | 5.68% (458) |

---- = other trematodes not recorded
Years at intervention lakes after intervention are marked with gray shading

**Fig 1.**
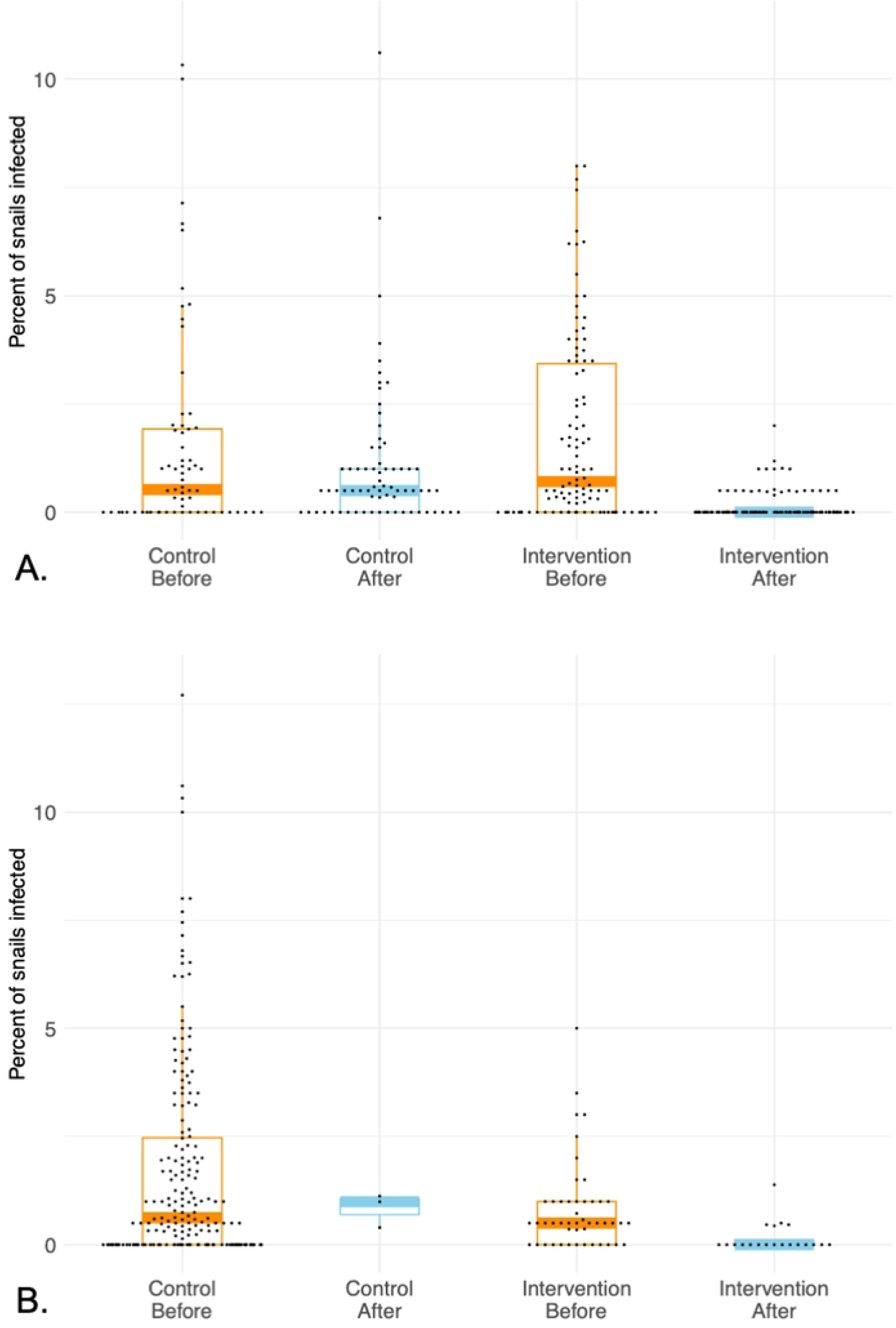
Snail infection data in BACI analyses at intervention lakes. (A) Higgins Lake as intervention lake. (B) Crystal Lake as intervention lake. Data are the percent of *Stagnicola emarginata* snails shedding schistosome cercariae with each dot representing a unique collection of snails. Boxes represent the first quartile to the third quartile, single vertical lines represent the positive 95% confidence interval, and thick horizontal lines represent the median. Orange color highlights time before intervention; blue color highlights time after intervention. Some snail collections did not have any snails shedding schistosome cercariae (0% snails infected), and where this outcome is frequent they appear as a solid line of zero values. Data represent 58996 snails examined in this study plus applicable historical data from Table 2.

**Fig 2.**
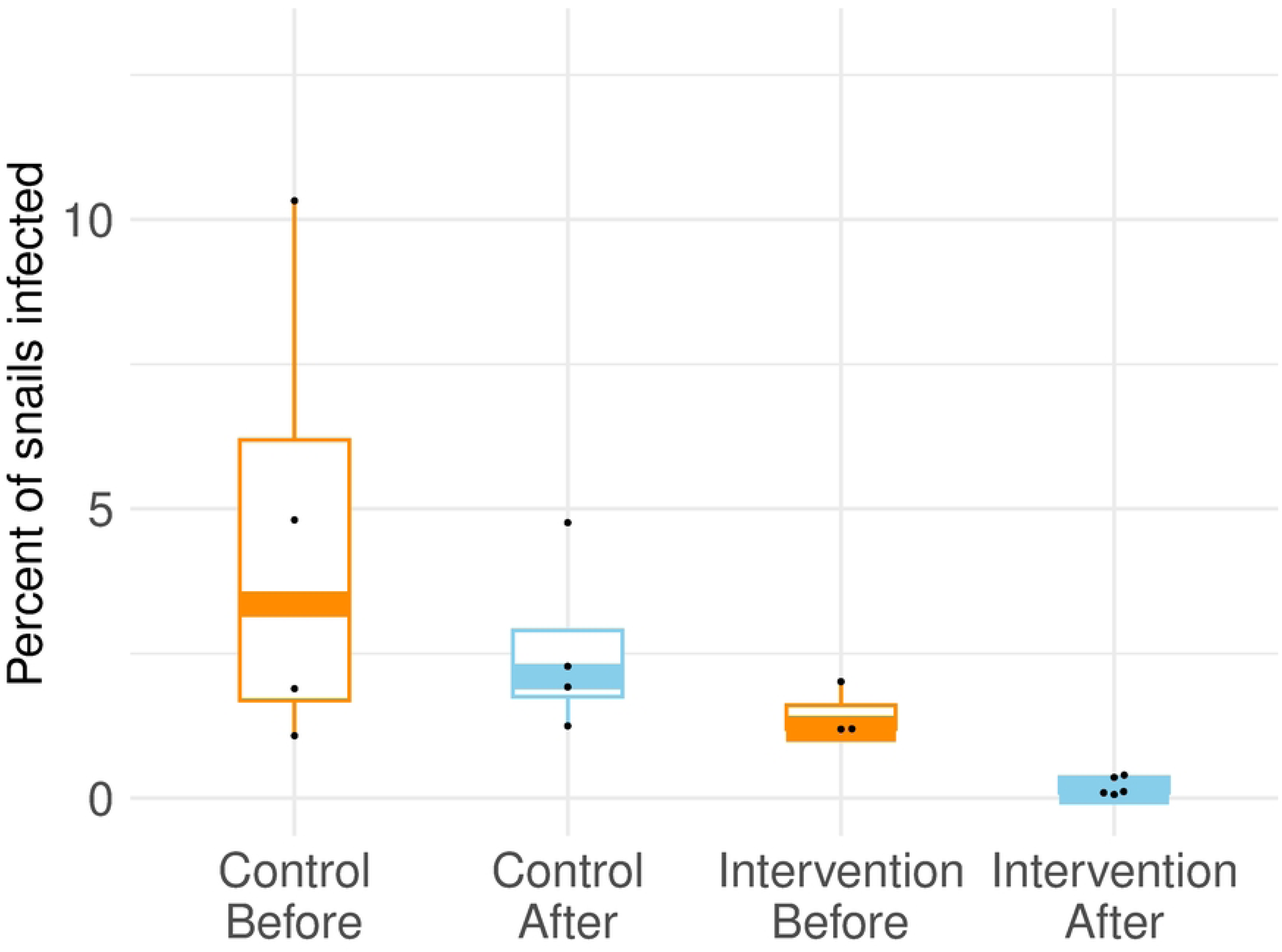
Historical snail infection data used in BACI analysis of Big Glen Lake. Data are the percent of *Stagnicola emarginata* snails shedding schistosome cercariae with each dot representing a survey report in the literature [18,22,25,44,45]. Boxes represent the first quartile to the third quartile, single vertical lines represent the positive 95% confidence interval, and thick horizontal lines represent the median. Orange color highlights time before intervention; blue color highlights time after intervention. Data represent a total of 123835 snails examined.

**Table 4.** Percent of *S. emarginata* snails shedding schistosome cercariae in mid-July collections at ten sites each on Higgins Lake and Crystal Lake, before and after intervention.

| Collection site | Higgins Lake |  |  |  |  |  | Crystal lake |  |  |
| --- | --- | --- | --- | --- | --- | --- | --- | --- | --- |
|  | 2015 | 2016 | 2017 | 2019 | 2020 |  | 2016 | 2018 | 2020 |
| 1 | 4.20% | 0.00% | 0.00% | 0.00% | 0.00% |  | 1.00% | 0.00% | 0.00% |
| 2 | 3.50% | 0.00% | 0.50% | 0.00% | 0.00% |  | 2.50% | 0.00% | 0.00% |
| 3 | 0.50% | 0.00% | 0.00% | 0.00% | 0.00% |  | 1.00% | 0.00% | 0.00% |
| 4 | 2.00% | 0.50% | 0.00% | 0.48% | 0.00% |  | 0.50% | 0.46% | 0.00% |
| 5 | 5.50% | 0.50% | 0.50% | 0.00% | 0.00% |  | 3.50% | 0.46% | 0.00% |
| 6 | 1.90% | 0.00% | 0.00% | 0.00% | 0.00% |  | 0.00% | 0.00% | 0.00% |
| 7 | 6.20% | 1.00% | 0.40% | 0.00% | 0.00% |  | 1.00% | 1.38% | 0.00% |
| 8 | 2.50% | 0.00% | 0.00% | 0.00% | 0.00% |  | 0.00% | 0.00% | 0.00% |
| 9 | 0.50% | 1.00% | 0.00% | 0.00% | 0.00% |  | 0.50% | 0.00% | 0.00% |
| 10 | 3.50% | 2.00% | 0.00% | 0.00% | 0.00% |  | 0.50% | 0.50% | 0.44% |
| Lake-wide total | 2.90% | 0.50% | 0.14% | 0.07% | 0.00% |  | 1.05% | 0.28% | 0.05% |
|  | n = 1757 | n = 2000 | n = 2185 | n = 1383 | n = 2063 |  | n = 2000 | n = 2112 | n = 2211 |

Shades of red indicate values greater than 0.25%.
Shades of blue indicate values less than 0.25%.
Gray shading indicates years after intervention by brood relocation.

The lake-wide percentage of all non-schistosome trematodes varied among lakes and years from 3.0% to 9.8% (Table 3). Lake-wide percentages of snails infected with five trematode families are visualized in Fig 3. Schistosomatidae was the only family that decreased substantially on both intervention lakes and remained low following merganser relocation (Fig 3).

**Fig 3.**
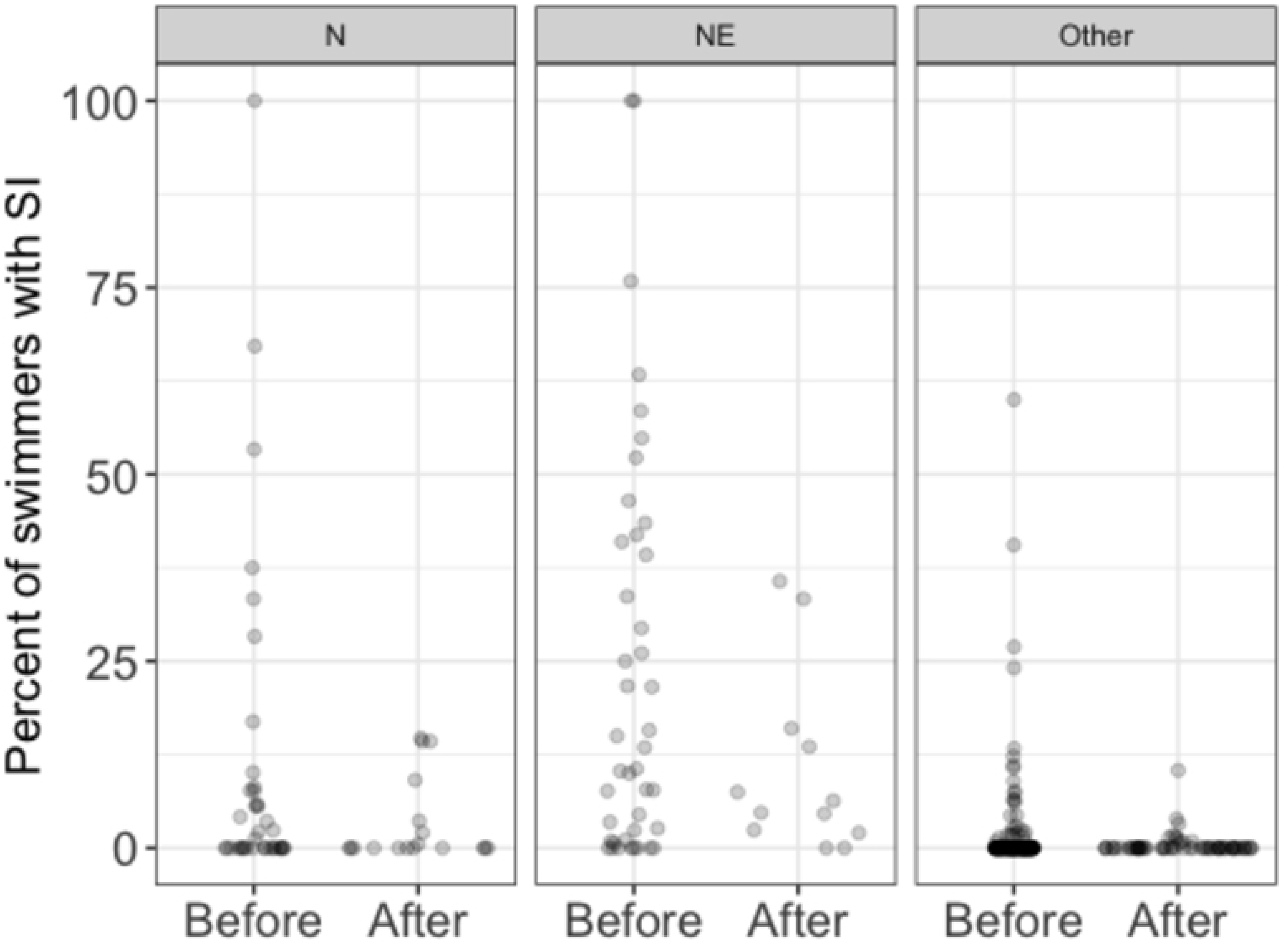
Percentages of *S. emarginata* snails shedding cercariae of five trematode families at intervention lakes. (A) Higgins Lake. (B) Crystal Lake. Schistosomatidae is the only trematode family that experienced a sustained decrease. Years of sample collection are depicted on x-axis (snails were not collected in 2018 at Higgins Lake or in 2017 and 2019 at Crystal Lake). Data represent 44021 snails examined.

### Crystal Lake CSA beach data

Onshore NNE winds were highly predictive of swimmer’s itch in the logistic regression model of CSA beach data (as expected, X^2^=115.87, p < 2.2 x 10^-16^), and swimmer’s itch cases per swimmer were substantially lower in the years after common merganser relocation (Fig 3, X^2^= 20.06, p = 7.5 x 10^-6^), regardless of wind direction (Fig 4). The percent of swimmers with SI ranged from 2.9% to 5.3% in 2013-2017 but fell to 1.7% in 2018 and 0.6% in 2019 (Fig 5). The model predicts a steep drop in SI rates during onshore NNE wind conditions, from 14.4% to 3.3%. Rates of SI are much lower in all other wind conditions, and the predicted reduction is correspondingly smaller, from 0.9% to 0.3%.

**Fig 4.**
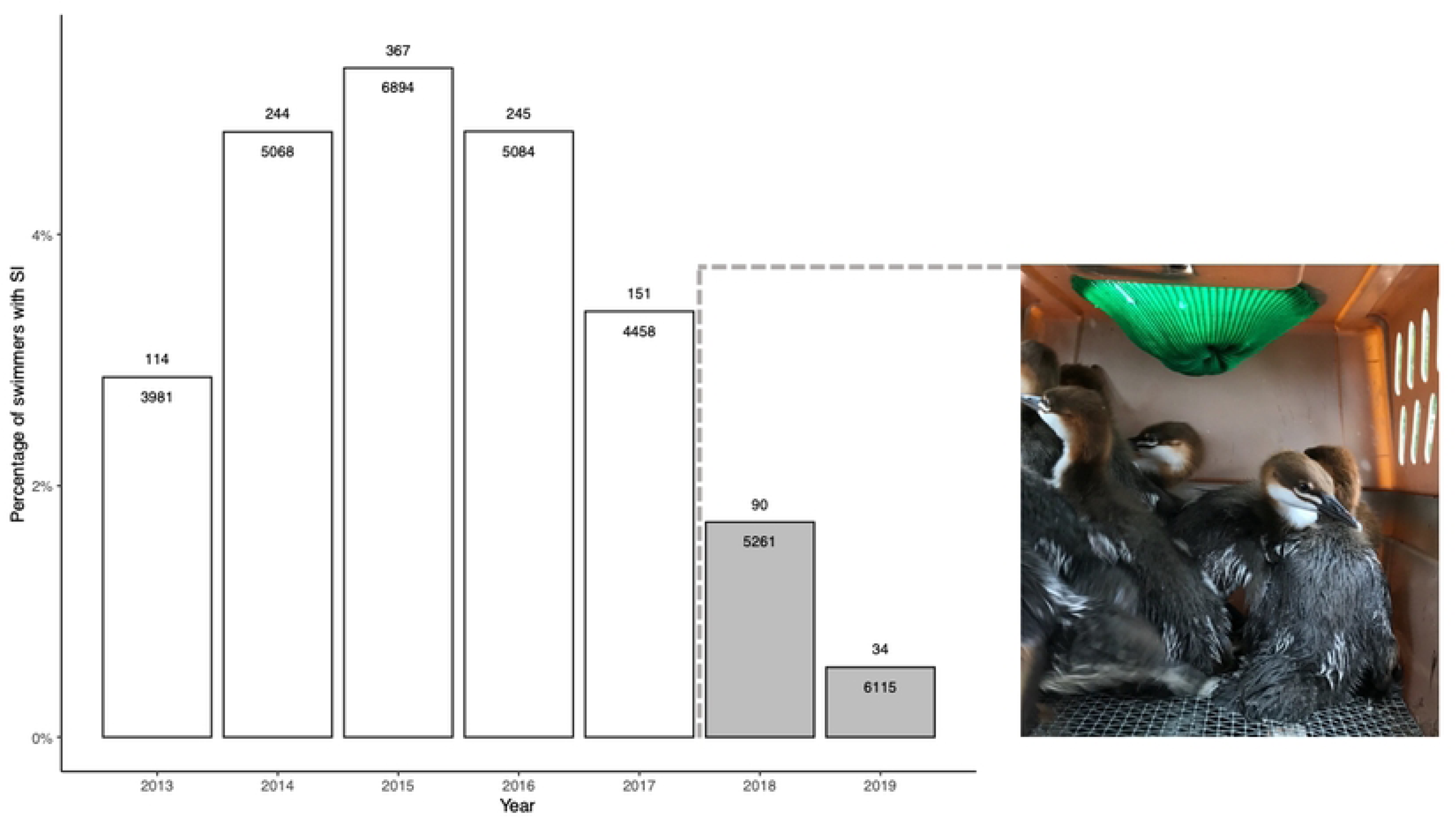
Percent of daily swimmers reporting swimmer’s itch at CSA beach before (2013-2017) and after (2018-2019) common merganser relocation. Data aggregated by three wind conditions: onshore N winds, onshore NE winds, and all other wind directions. Each dot represents a day. Since many days did not have swimmer’s itch cases, especially without onshore N or NE winds, they appear as overlapping zero values at the bottom of the plots.

**Fig 5.**
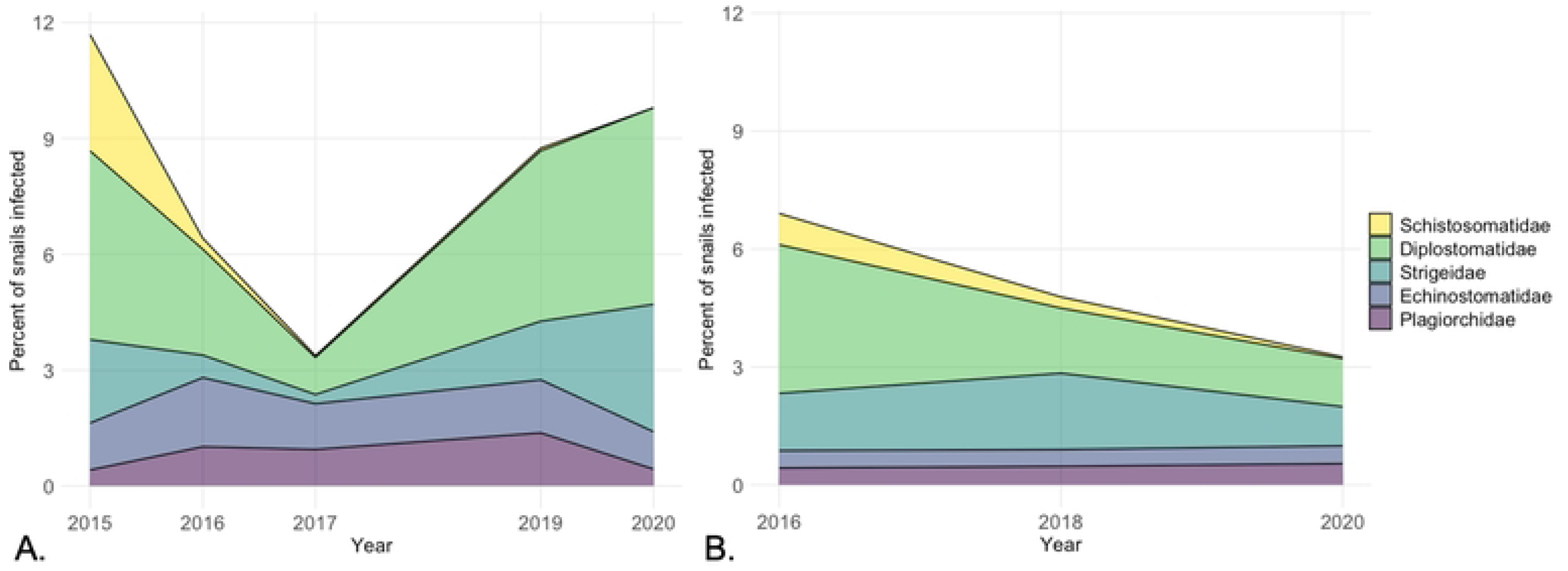
Percent of swimmers reporting swimmer’s itch annually at CSA beach, Crystal Lake. Number of swimmer’s itch cases above and total number of swimmers below the top of each column. Dashed line, photo, and grayed columns highlight years after common merganser relocation.

## Discussion

### Schistosome population reduction at Higgins Lake and Crystal Lake

Annual common merganser brood relocation had profound effects on the lake-wide percentage of snails infected with *T. stagnicolae* at both intervention lakes (Fig 1, Tables 3,4). All years after relocation began were substantially lower than years before relocation. The effects were immediate, with strong reductions in the first year on both Higgins (10-fold) and Crystal Lakes (3-fold). Reductions were sustained over multiple years and reached similarly low levels (∼0.05% or lower) on both lakes, a greater than 50-fold total reduction on Higgins Lake and greater than 10-fold total reduction on Crystal Lake. It is important to note that these reductions in schistosome populations occurred despite the yearly presence of 2^nd^-year common mergansers during the summer as well as much larger migrant populations in spring and fall on both lakes. Lethal take of adult birds by Gerrish township officials unassociated with us in spring 2015 and 2016 only removed a fraction of the migratory population on Higgins Lake and our limited lethal take for other scientific purposes left more adults at both lakes in the summer than it eliminated.. Therefore, only relocation of common merganser broods can account for the immediate, severe, and sustained decline in snail infection prevalence.

The BACI-design of the study ensures that the reduction in snails infected with schistosomes is not due to pre-existing differences among lakes, a regional decrease in *T. stagnicolae* populations that affects all lakes, or an artifact of the timing of sampling at different lakes. Furthermore, our records of other trematode infections strongly suggest that our assessment tool of shedding snails remained effective throughout the study, and importantly, that schistosomes were the only trematode group that showed sustained reduction after common merganser brood relocation (Fig 2). Finally, the data from intervention lakes before intervention, the data from control lakes, and the historical data demonstrate that under natural conditions, *T. stagnicolae* population levels in snails vary from year to year but do not typically decline to the low levels seen in this study following common merganser brood relocation (Tables 3,4).

### SI reduction at Crystal Lake

A reduction of SI cases in humans is the ultimate objective of any SI mitigation effort. The data from CSA beach on Crystal Lake is a 7-year data set of almost 37,000 swimmer sessions that provides an excellent measure of whether the decrease in snail infection rates translates into fewer SI cases. The number of SI cases at CSA beach dropped significantly in the two years following common merganser brood relocation, correspondent to reduced snail infection (Figs 3 and 4). To our knowledge, this is the first time that SI reduction has been documented with case data controlled for the number of swimmers entering the water and includes years prior to and after mitigation efforts have begun. In addition, while the CSA study did not include the number of papules per swimmer in the study design, the study supervisors anecdotally reported to us that a reduction in the severity of the cases (in terms of the number of papules) was frequently expressed by the lifeguards in 2018 and 2019 (Leslie Ritter, Al Flory, personal communication). Finally, while snail density is one factor that could influence the number of SI cases, this did not seem to be a factor at CSA beach. Snail densities at CSA beach were relatively low in years prior to [46] and after (unpublished data) common merganser brood relocation.

### Schistosome population reduction at Big Glen Lake

A similar mitigation program was performed in the 1980’s at Big Glen Lake in northern Michigan [25]. That study also reported immediate and sustained reductions in the percentage of snails infected, up to 6 years after mitigation efforts began, and reductions in snail infection percentages are remarkably similar to those achieved on Crystal Lake and Higgins Lake (Fig 1, Tables 1 and 3). This mitigation program also targeted the common merganser definitive host, though efforts involved a mixture of relocation of broods, praziquantel treatment of trapped broods with re-release on Big Glen Lake (no relocation), and lethal take. The data were not analyzed statistically in the original study [25], but in our reanalysis here we found strong statistical inference that the percentage of snails infected was substantially lower because of the mitigation efforts, making this a third instance of a definitive host focused approach being successful in lowering the whole-lake population of *T. stagnicolae* by more than an order of magnitude.

### Factors in success

A number of biological factors contribute to the success of mitigation on Crystal Lake, Higgins Lake, and Big Glen Lake: 1) *S. emarginata* is the most abundant snail on these three lakes (personal observations, [28]) 2) *T. stagnicolae* is the only known avian schistosome in the area that uses *S. emarginata* snails; and 3) common merganser broods, the only known hosts for *T. stagnicolae,* are as abundant or more abundant than other waterfowl broods on these lakes. These three factors combine to make *T. stagnicolae* the dominant parasite on these lakes and other lakes like them in northern Michigan, as noted in historical studies [19–21] and confirmed by recent ones [17,18]. In addition, as is true for many pulmonate snails in temperate regions, adult *S. emarginata* populations decline in late summer and are replaced by young snails that hatched earlier in the summer, requiring *T. stagnicolae* to infect a new cohort of snails each year. Relocation of summer broods reduces transmission to the next generation of snails.

### Factors for further investigation

In addition, other biological aspects of the *T. stagnicolae* life cycle may have contributed and merit further investigation. First, the importance of common merganser ducklings in propagating *T. stagnicolae* is strongly supported by our data, but much more could be done to determine why that is so. If ducklings release more parasite eggs on a consistent basis than adults, how long does this occur, and is it because ducklings are initially infected with many adult worms and some worms die as the ducklings mature, or is it because most worm pairs survive but become more limited in egg production by a maturing avian immune system? Are eggs and/or miracidia from ducklings perhaps more viable or infective to snails than those from adults (see [49])? Second, how important is the co-incidence of ducklings with young naïve snails in the early summer? We suspect that younger *S. emarginata* snails are more susceptible to *T. stagnicolae*, as has been found for other avian, mammalian, and human schistosomes: *Trichobilharzia szidati* in *Lymnaea stagnalis* [50,51]*, Schistosomatium douthitti* in *Stagnicola emarginata* [52], and *Schistosoma mansoni* in *Biomphalaria* [53–55], though the mechanisms responsible for this age-related effect vary among these parasite-host systems. Finally, while mitigation succeeded in severely reducing the population of *T. stagnicolae* on experimental lakes, even with 100% of broods relocated from the system the parasite is not eliminated, likely due to minor contributions from fall and spring migrants, prospecting breeding pairs in the spring, and 2^nd^-year non-breeding adults that can be present in the summer months. Which of these (migrants, prospecting pairs, or 2^nd^-year adults) is of next-tier importance in maintaining the parasite’s population on a lake? If migrants contribute, are fall or spring migrants more important and is transmission more limited by cold water temperatures or by snail age, or both?

### Contrasts with a recent study

Our data contradict the study outcomes of Rudko et al. [28], who concluded that common merganser relocation on Big Glen Lake and North Lake Leelanau in northern Michigan in 2017-2019 did not meaningfully impact *T. stagnicolae* populations, and that migrating common mergansers were sufficient to sustain *T. stagnicolae* populations. The difference in conclusion may be due to several factors, including differences in study design. Rudko et al. [28] did not present data from before relocation on their study lakes and could only make comparisons between control and intervention lakes (a CI or Control-Intervention study), which has less power than a BACI to detect differences related to an intervention because prior differences between the lakes will not be accounted for [33,34]. In addition, there were challenges in successfully trapping common merganser broods in a timely manner which may have decreased the effectiveness of the program on those lakes (personal communication with one of the trappers, and also documented at https://www.glenlakeassociation.org/swimmers-itch-control-update/ [56]). The Big Glen Lake and North Lake Leelanau programs also assumed that broods would not pass parasite eggs until four weeks of age [56]. However, in the laboratory, Peking ducks (a breed of the mallard *Anas platyrynchos*) and black ducks (*Anas rubripes*) can be passing *T. szidati* eggs at 13-14 days after infection with peak egg passage occurring at 15-22 days post infection [49,57]. Common merganser broods at Crystal Lake and Higgins Lake were usually relocated at about 2 weeks of age (or earlier), before most worms could mature and this likely contributed to the strategy’s success on those two lakes.

We assessed *T. stagnicolae* populations by a simple and long-standing technique: the frequency of patent infections among large numbers of collected snails [12,58]. We view the percentage of infected snails as the most direct measure of the population levels of the parasite on a lake, because it is directly related to the amount of successful transmission from common mergansers to snails occurring over the previous year. While the cercarial shedding technique can potentially underestimate infection prevalence due to variable shedding response or immature infections, it will not be systematically biased relative to whether common merganser relocation has or has not occurred. Moreover, the common presence of other trematode infections suggests that our shedding methods were consistently effective and underestimation of schistosome infections was minimal.

In contrast, Rudko et al. [28] utilized qPCR to estimate the number of cercariae in 25 L surface water samples taken weekly from multiple sites on each lake and utilized a citizen science network for sample collection. Advantages of this approach include the ability to sample more sites more frequently and citizen scientists can be trained to properly collect samples [12,58,59]. However, this measure is multiple steps removed from the number of snails infected and the amount of merganser to snail transmission since each snail infected does not produce the same number of cercariae. In fact, the number of cercariae a given snail produces into the water varies day to day even in laboratory studies [60,61]. Moreover, representative sampling of such large bodies of water is difficult and undoubtedly affected by daily variances in environmental conditions because cercariae can be dispersed or concentrated by wind and water currents. Finally, whether qPCR data accurately estimates the number of live cercariae in the sample is further complicated by the ubiquity of schistosome environmental DNA (eDNA) in the water due to the daily release and death of cercariae, and the poorly understood dynamics of eDNA stabilization and decomposition [62–64]. That eDNA is frequently used today to detect rare organisms using small (1-2 L) sample volumes [65] should caution against methods which assume that all DNA extracted from a much larger sample (25 L) is from live cercariae directly collected in the sample.

### Applications and areas of future research

We conclude that the program of common merganser brood relocation has been remarkably successful at Crystal Lake and Higgins Lake. Results from these two geographically separated lakes, as well as similar past results at Big Glen Lake, demonstrate that mitigation efforts targeting the definitive hosts can be very effective at reducing the number of snails infected and consequently, the number and severity of SI cases, while avoiding the use of copper sulfate or other chemicals. Additional benefits of the brood relocation strategy at these lakes include 1) elimination of incentive for lakefront owners and businesses to shoot common mergansers, legally or illegally (we have witnessed legal hunting in the migratory season as well as illegal hunting in the breeding season on multiple lakes in Michigan), and 2) non-breeding adult and migratory populations can be left undisturbed.

Relocated common merganser broods appear to do well (personal observations), an unsurprising outcome since they were relocated to locations where this species naturally breeds. Although we identified above some features that were unique to the lake systems studied here, our results suggest efforts that target adult schistosomes in the vertebrate host, especially in the young-of-the-year birds, can have a strong effect.

Lessons from this approach are likely adaptable to other systems where different avian schistosome species and different waterfowl hosts are involved, but many factors need to be considered on a lake-by-lake basis and the strategy will not be viable in all situations. Thorough investigation should be conducted into which parasite species and corresponding snail and vertebrate hosts are contributing to a particular SI problem, and adjustments will have to be made for each unique system so that efforts are correctly targeted. In our study systems *T. stagnicolae* appears to be adapted to take advantage of the appearance of ducklings to achieve high population levels, and we predict that at least some avian schistosomes are likely to share this strategy. However, we also emphasize that other avian schistosomes could differ in degree of reliance on young-of-the-year birds.

Another factor that will vary in each instance is feasibility of host relocation, due to the lack of a nearby suitable relocation site, the number of waterfowl involved, the size of the waterbody, or species-specific restrictions on such activity. We do not advocate lethal take for SI control purposes, so in these cases, praziquantel, a highly effective therapy against schistosomes and other parasites, might be used to reduce the parasite egg output of broods during the key summer months. Yet whether praziquantel alone can be as effective in whole-lake reduction of snail infections and SI cases needs to be studied. A disadvantage is that praziquantel impacts only the adult worms already present and does not prevent new infections, so young-of-the-year would likely need to be treated at least twice to prevent them from passing parasite eggs during the summer months [25,66,67]. Fortunately, praziquantel has very low toxicity for vertebrates, is metabolized quickly, and is tolerated well, hence its widespread and heavy use in the treatment of parasitic helminths in pets, fish farming, captive and free wildlife, livestock, and humans [68,69]. Praziquantel is often successful in treating helminth infections in birds without known side effects [70–72], including the treatment of schistosomes in waterfowl [25,26,67,73]. A few authors suggest continued research of derivatives/metabolites and their possible effects on invertebrates in the environment [69,74], but the small amounts needed and the diluting effect of large lakes should minimize this concern. Experts in ecotoxicology and veterinary science should be consulted to determine appropriate doses, mode of delivery, expected degree of environmental dilution and degradation of excreted drug compounds, and any environmental risks. Finally, simple and inexpensive actions that may reduce waterfowl host populations should also be considered, such as encouraging the public not to feed waterfowl, removing or blocking nest sites before they are used, promoting natural shoreline vegetation that can discourage lawn grazers like mallards and Canada goose, and minimizing roosting sites for young broods along the shore.

One interesting application of this study may be to SI in a much different context: that of rice paddies in central and eastern Asia, where there has been increasing recognition that field workers experience high rates of SI [75–78]. In many places, especially where organic practices prohibit the use of pesticides, large flocks of domesticated ducks are released to roam flooded fields to consume insects and other pests. Schistosome infections are known in these domestic flocks and are likely leading to more snail infections and increased worker exposure. Perhaps duck flocks can be periodically treated with praziquantel to rid them of worms before times of release and again later to purge any new infections that may have been picked up. Knowing when young snails appear and whether they are significantly more susceptible could potentially enhance effectiveness of a praziquantel-based strategy.

The impressive reduction of an avian schistosome population in our study serves as a reminder to efforts combatting human schistosomiasis that any interventions or practices that decrease contact between human waste and water sources can potentially reduce transmission to snails in the environment, which will in turn reduce transmission risk of new human infections [79–82]. Although there have been recent calls to re-integrate snail control with the ongoing mass administration of praziquantel [83,84], there also remains an inadequate emphasis on structural changes that affect human interactions with water [82]. A notable difference between SI and human schistosomiasis is that human schistosome transmission can require year-round vigilance because it occurs in tropical and subtropical environments, whereas the success of the Michigan strategy benefits from the seasonality of transmission, and mitigation efforts can focus on a relatively narrow time frame (∼2 summer months).

With the success of the vertebrate host-focused strategy described here there is also still much to learn. Since relocation and praziquantel based approaches are labor-and cost-intensive, perhaps our findings will inspire alternative vertebrate host focused strategies. In places where relocation has succeeded, there are also questions about how best to use resources once SI has been reduced. Could relocation or praziquantel programs be implemented on an alternate year basis with only a minimal and tolerable increase in SI? Without common merganser brood relocation, will snail infection percentages return to pre-intervention levels and how quickly? Other questions surround what should be defined as success. The low lake-wide snail infection percentages and decrease in SI cases reported here have satisfied lake residents and associations who sought to reduce a public health and economic problem. What is a tolerable lake-wide snail infection percentage, and might it differ among lakes, interacting with lake attributes like lake topography and the size and density of snail populations? The public health, recreational, and economic benefits should be continually evaluated relative to any potential ecological costs and considered alongside the costs and benefits of alternative strategies if they are developed.

## Conclusion

Here we have demonstrated that a strategy focused on young-of-the-year vertebrate hosts can significantly lower the population of a swimmer’s itch-causing avian schistosome and reduce SI cases, avoiding the use of environmental toxins like copper sulfate. The dramatic and rapid results suggest that the parasite studied, *T. stagnicolae*, strongly relies on young-of-the-year common mergansers to achieve high population levels on lakes in northern Michigan, and that relocation must be timely to achieve success. Applicability of the strategy to other SI-causing schistosomes may differ according to the schistosome species biology.

## Supporting information

**S1 Fig. Percentage of snails infected is highest from late June to late July, based on temporal data in three lakes.** Higgins 2016 and 2017 collections (in lighter shades of blue) are after mitigation had begun and highest levels were still detected in mid-July. Consequently, all later collections (2018-2020) occurred in mid-July. Data represent 44310 snails examined.

## Acknowledgements

We recognize Ronald L. Reimink and his pioneering research and work with HDB to advance knowledge about swimmer’s itch and the role of common mergansers in parts of Michigan. RLR also contributed to this study in 2015 and 2016, as did Chris Froelich and Kelsey Froelich, by collecting all snail data at Glen Lake, North Lake Leelanau, and Crystal Lake in 2015-2016 and 50% of the snail data at Higgins Lake in 2015-2016. We thank Ren Tubergen, Emily Blankespoor, Bradley Scholten, Claire DeJong, and Elias DeJong for helping the authors collect snail data in 2017-2020. We are especially grateful to Leslie Ritter, the late Al Flory, Dave Wynne, and CSA staff for collecting and sharing CSA beach data. We thank the University of Michigan Biological Station and especially its staff for excellent logistic support. Thanks to Stacy DeRuiter for providing statistical advice. Special thanks to Bob and Diane Wagner and many other lake residents and volunteers who helped in so many ways. This study would never have happened without the visionary community leadership of Ken Dennings, Jim Vondale, and the Higgins Lake Swimmer’s Itch Organization (HLSIO) now led by Greg Semack.

